## Supplementary Table 1 for "*De novo* fatty-acid synthesis protects invariant NKT cells from cell death, thereby promoting their homeostasis and pathogenic roles in airway hyperresponsiveness"

**Table S1. List of reagents used for flow cytometry**

| Antibody/Reagent | Catalogue # | Clone | Company |
|---|---|---|---|
| PerCP/Cyanine5.5 anti-mouse CD45 | 103132 | 30-F11 | BioLegend |
| PE/Cy7 anti-mouse CD11c | 117318 | N418 | BioLegend |
| APC anti-mouse F4/80 | 123116 | BM8 | BioLegend |
| PE anti-mouse/human CD11b | 101208 | M1/70 | BioLegend |
| FITC anti-mouse F4/80 | 123108 | BM8 | BioLegend |
| PE anti-mouse CD170 (Siglec-F) | 155506 | S17007L | BioLegend |
| Zombie Aqua™ Fixable Viability Kit | 423101 | | BioLegend |
| Brilliant Violet 421™ anti-mouse/human CD11b | 101236 | M1/70 | BioLegend |
| PE – Annexin V | 640947 | - | BioLegend |
| PE- anti-mouse PLZF | 145803 | Mag.21F7 | BioLegend |
| PBS 57 – APC conjugated anti-mouse CD1d tetramer | - | - | NIH Tetramer Facility |
| PerCP anti-mouse CD45.1 | 110726 | A20 | BioLegend |
| Alexa Fluor® 700 anti-mouse CD45.2 | 109822 | 104 | BioLegend |
| PE/Cy7 anti-mouse TCR β chain | 109222 | H57-597 | BioLegend |
| APC anti-mouse CD86 | 105012 | GL-1 | BioLegend |
| FITC anti-mouse CD80 | 104706 | 16-10A1 | BioLegend |
| PE/Cy7 anti-mouse CD69 | 104511 | H1.2F3 | BioLegend |
| FITC anti-mouse GLUT1 | Sc-377228 | A-4 | Santa Cruz |
| FITC anti-mouse Ly-6G | 127606 | 1A8 | BioLegend |
| PE anti-mouse CD19 | 152408 | 1D3/CD19 | BioLegend |
| APC anti-mouse CD4 | 100516 | RM4-5 | BioLegend |
| FITC anti-mouse CD8a | 100706 | 53-6.7 | BioLegend |
| PE anti-mouse FOXP3 | 126404 | MF-14 | BioLegend |
| Ovalbumin, Fluorescein Conjugate | O23020 | | ThermoFisher Scientific |
| PerCP anti-mouse NK-1.1 | 108725 | PK136 | BioLegend |
| APC anti-mouse/human CD44 | 103011 | IM7 | BioLegend |
| FITC anti-mouse BrdU | 347583 | B44 | BD |
| PE anti-mouse CD24 | 553261 | M1/69 | BD Bioscience |
| Brilliant Violet 421™ anti-mouse CD62L | 104435 | MEL-14 | BioLegend |
| PE anti-mouse CD64 (FCγRI) | 139303 | X54-5/7.1 | BioLegend |
| Brilliant Violet 421™ anti-mouse Ly-6C | 128031 | HK1.4 | BioLegend |
| PE/Cy7 Armenian Hamster IgG Isotype Ctrl | 400921 | HTK888 | BioLegend |
| FITC Rat IgG2a, κ Isotype Ctrl | 400505 | RTK2758 | BioLegend |
| PE Rat IgG2a, κ Isotype Ctrl | 400507 | RTK2758 | BioLegend |
| Brilliant Violet 421™ Rat IgG2b, κ Isotype Ctrl | 400639 | RTK4530 | BioLegend |
| PerCP/Cyanine5.5 anti-mouse IL-4 | 504123 | 11B11 | BioLegend |
| PE anti-mouse IL-13 | 159403 | W17010B | BioLegend |
| APC anti-human CD25 | 302609 | BC96 | BioLegend |
| FITC anti-human CD4 | 317407 | OKT4 | BioLegend |
