## Supplementary Table 2 for "*De novo* fatty-acid synthesis protects invariant NKT cells from cell death, thereby promoting their homeostasis and pathogenic roles in airway hyperresponsiveness"

| Species | Gene | F/R | Sequence |
|---|---|---|---|
| Mouse | Acox1 | F | CTG CCA AGG GAC TCC AGA GCA GCT |
| Mouse | Acox1 | R | GAC ATG GAC ACA TCC ACC ATG CAG |
| Mouse | Fasn | F | AGC GGC CAT TTC CAT TGC CC |
| Mouse | Fasn | R | CCA TGC CCA GAG GGT GGT TG |
| Mouse | Acc1 | F | ACA GTG GAG CTA GAA TTG GAC |
| Mouse | Acc1 | R | ACT TCC CGA CCA AGG ACT TTG |
| Mouse | Ppar-gamma | F | GTG ATG GAA GAC CAC TCG CAT T |
| Mouse | Ppar-gamma | R | CCA TGA GGG AGT TAG AAG GTT C |
| Mouse | Tbx21 | F | TTCCCATTCCTGTCCTTCAC |
| Mouse | Tbx21 | R | CCACATCCACAAACATCCTG |
| Mouse | Gata3 | F | GGAAACTCCGTCAGGGCTA |
| Mouse | Gata3 | R | AGAGATCCGTGCAGCAGAG |
| Mouse | Rorc | F | TGA GGC CAT TCA GTA TGT GG |
| Mouse | Rorc | R | CTT CCA TTG CTC CTG CTT TC |
| Mouse | Foxp3 | F | CCC AGG AAA GAC AGC AAC CTT |
| Mouse | Foxp3 | R | TTC TCA CAA CCA GGC CAC TTG |
| Mouse | Hk2 | F | AGA GAA CAA GGG CGA GGA G |
| Mouse | Hk2 | R | GGA AGC GGA CAT CAC AAT C |
| Mouse | g6pase | F | CCA TGC AAA GGA CTA GGA ACA A |
| Mouse | g6pase | R | TAC CAG GGC CGA TGT CAA C |
| Mouse | Fbp1 | F | CCA TCA TAA TCG AAC CTG AG |
| Mouse | Fbp1 | R | CTT CTC AGA AGG CTC ATC AG |
| Mouse | Sdhb | F | CTA AAT AAG TGC GGA CCT ATG G |
| Mouse | Sdhb | R | AGT ATT GCC TCC GTT GAT GTT C |
| Mouse | Cpt1a | F | CCA TCC TGT CCT GAC AAG GTT TAG |
| Mouse | Cpt1a | R | CCT CAC TTC TGT TAC AGC TAG CAC |
| Mouse | Pkm2 | F | CTG GCT CAG AAG ATG ATG ATC G |
| Mouse | Pkm2 | R | CTT GGT GAG CAC GAT AAT GG |
| Mouse | cEbpa | F | CAA AGC CAA GAA GTC GGT GGA |
| Mouse | cEbpa | R | TCA TTG TGA CTG GTC AAC TCC AGC |
| Mouse | Bak1 | F | ATA TTA ACC GGC GCT ACG AC |
| Mouse | Bak1 | R | AGG CGA TCT TGG TGA AGA GT |
| Mouse | Bax | F | TAG CAA ACT GGT GCT CAA |
| Mouse | Bax | R | TCT TGG ATC CAG ACA AGC AG |
| Mouse | Bcl-2 | F | CTC GTC GCT ACC GTC GTG ACT TCG |
| Mouse | Bcl-2 | R | CAG ATG CCG GTT CAG GTA CTC AGT C |
| Mouse | Bcl-XL | F | TGG AGT AAA CTG GGG GTC GCA TCG |
| Mouse | Bcl-XL | R | AGC CAC CGT CAT GCC CGT CAG G |
| Mouse | Gapdh | F | GGGAAGCTCACTGGCATGG |
| Mouse | Gapdh | R | CTTCTTGATGTCATCATACTTGGCAG |
| Mouse | IFN-r | F | CGG CAC AGT CAT TGA AAG CCT A |
| Mouse | IFN-r | R | GTT GCT GAT GGC CTG ATT GTC |
| Mouse | IL-4 | F | TCA ACC CCC AGC TAG TTG TC |
| Mouse | IL-4 | R | TGT TCT TCG TTG CTG TGA GG |
| Mouse | IL-5 | F | CTC TGT TGA CAA GCA ATG AGA CG |
| Mouse | IL-5 | R | TCT TCA GTA TGT CTA GCC CCT G |
| Mouse | IL-13 | F | CCT GGC TCT TGC TTG CCT T |

| | | | |
|---|---|---|---|
| Mouse | IL-13 | R | GGT CTT GTG TGA TGT TGC TCA |
| Mouse | IL-17a | F | GGT CTT GTG TGA TGT TGC TCA |
| Mouse | IL-17a | R | GGG TCT TCA TTG CGG TGG AGA G |
| Mouse | Fapb1 | F | AGG AGT GCG AAC TGG AGA CCA T |
| Mouse | Fapb1 | R | GTC TCC ATT GAG TTC AGT CAC GG |
| Mouse | Fapb3 | F | AGA GTT CGA CGA GGT GAC AGC A |
| Mouse | Fapb3 | R | TTG TCT CCT GCC CGT TCC ACT T |
| Mouse | Fapb5 | F | GAC GAC TGT GTT CTC TTG TAA CC |
| Mouse | Fapb5 | R | TGT TAT CGT GCT CTC CTT CCC G |
| Mouse | Cxcr3 | F | CAG CCT GAA CTT TGA CAG AAC CT |
| Mouse | Cxcr3 | R | GCA GCC CCA GCA AGA AGA |
| Mouse | Cxcr4 | F | TCA GCC TGG ACC GGT ACC T |
| Mouse | Cxcr4 | R | GCA GTT TCC TTG GCC TTT GA |
| Mouse | Ccr4 | F | GGA CTA GGT CTG TGC AAG ATC G |
| Mouse | Ccr4 | R | TGC CTT CAA GGA GAA TAC CGC G |
| Mouse | Ccr6 | F | ACA GAG CCA TCC GAG TCG TGA T |
| Mouse | Ccr6 | R | CTG GTG TAG GCG AGG ACT TTC T |
| Human | PPARg | F | AGT CCT CAC AGC TGT TTG CCA AGC |
| Human | PPARg | R | GAG CGG GTG AAG ACT CAT GTC TGT C |
| Human | PPARg | F | AGC CTG CGA AAG CCT TTT GGT G |
| Human | PPARg | R | GGC TTC ACA TTC AGC AAA CCT GG |
| Human | Beta Actin | F | TCC CTG GAG AAG AGC TAC GA |
| Human | Beta Actin | R | AGC ACT GTG TTG GCG TAC AG |
| Human | ACC1 | F | ATG GGC GGA ATG GTC TCT TTC |
| Human | ACC1 | R | TGG GGA CCT TGT CTT CAT CAT |
| Human | FASN | F | GGA GGT GGT GAT AGC CGG TAT |
| Human | FASN | R | GGG TAA TCC ATA GAG CCC AG |
| Human | HK2 | F | GAG TTT GAC CTG GAT GTG GTT GC |
| Human | HK2 | R | CCT CCA TGT AGC AGG CAT TGC T |
| Human | IL4 | F | CCG TAA CAG ACA TCT TTG CTG CC |
| Human | IL4 | R | GAG TGT CCT TCT CAT GGT GGC T |
| Human | IL13 | F | ACG GTC ATT GCT CTC ACT TGC C |
| Human | IL13 | R | CTG TCA GGT TGA TGC TCC ATA CC |
| Human | IFNG | F | GAG TGT GGA GAC CAT CAA GGA AG |
| Human | IFNG | R | TGC TTT GCG TTG GAC ATT CAA GTC |
| Human | IL10 | F | TCT CCG AGA TGC CTT CAG CAG A |
| Human | IL10 | R | TCA GAC AAG GCT TGG CAA CCC A |
